# Establishment of an efficient transformation and CRISPR/Cas9-mediated gene editing system in Chinese local planting cassava (*Manihot esculenta* Crantz) cultivar SC8

**DOI:** 10.1101/2022.03.17.484691

**Authors:** Yajie Wang, Xiaohua Lu, Xinghou Zhen, Hui Yang, Yannian Che, Jingyi Hou, Ruimei Li, Jiao Liu, Mengting Geng, Xinwen Hu, Yuan Yao, Jianchun Guo

## Abstract

Cassava starch is a widely used raw material for industrial production. South Chinese cassava cultivar 8 (*Manihot esculenta* Crantz cv. SC8) is one of the main locally planted cultivars. In this study, an efficient transformation system for cassava SC8 mediated with *Agrobacterium* strain LBA4404 was presented for the first time, in which the factors of *Agrobacterium* strain cell infection (density OD600 = 0.65), 250 µM acetosyringone induction, and agro-cultivation with wet friable embryogenic callus (FEC) for 3 days in dark conditions were found to increase the transformation efficiency through the binary vector pCAMBIA1304 harboring GUS- and GFP-fused genes driven by the *CaMV35S* promoter. Based on the optimized transformation protocol, approximately 120-140 independent transgenic lines per mL settled FEC cell volume (SCV) by gene transformation in approximately five months, and 45.83% homozygous mono-allelic mutations of the *MePDS* gene with a *YAO* promoter-driven CRISPR/Cas9 system were generated. This study will open a more functional avenue for the genetic improvement of cassava SC8.

## 1. Introduction

Cassava (*Manihot esculenta* Crantz) is an important food crop in the tropics. The tuber roots of cassava are rich in starch, which is used as a staple food for 700 million people in 105 countries. Cassava provides food for humans and raw materials for industrial production, such as biofuel processing, paper products, starch processing, livestock feeds, the textile industry, and medical products (Parmar et al., 2017). Enhancing starch accumulation and modification of starch functional properties are important goals for cassava breeding. Due to cassava’s highly heterozygous genome and its low flowering and fruit setting rates (Souza et al., 2020; Ceballos et al., 2020), it takes longer to improve the characteristics of cassava cultivars by hybrid breeding technology.

Transgenic technology is considered to be a powerful tool for the genetic improvement of cassava. *Agrobacterium*-mediated gene transformation or editing in the model cultivar (*Manihot esculenta* cv TMS60444) has been used to modify cassava starch properties. For example, through downregulation of the granule-bound starch synthase gene, *GBSSI* expression in cv TMS60444 by RNAi technology, the amylose content in tuber roots of the transgenic cassava was significantly reduced (<5%) in comparison with that of the wild type (approximately 25%). The values of clarity, peak viscosity, gel breakdown, and swelling index were increased. In contrast, setback, consistency, and solubility were notably reduced (Zhao et al., 2011). Using gene editing technology to knock out the *GBSSI* gene could eliminate amylose starch content in cv TMS60444 tuber roots and lead to a lower peak temperature, higher peak viscosity, and higher final viscosity compared to those of WT in a recent study (Bull et al., 2018). Inhibition of the expression of starch branching enzyme gene *MeBE2* in cv TMS60444 by RNAi could increase high-amylose starch in cassava tuber roots, and this was increased by up to 50% and had a higher melting temperature in a study that used this method (Zhou et al., 2020). Overexpressing the potato *StGWD* gene or downregulating endogenous phosphoglucan phosphatase genes (*MeSEX4* or *MeLSF2*) could alter starch phosphorylation and starch properties in cv TMS60444 (Wang et al., 2018). However, all these studies were performed on model cassava cv TMS60444, which is amenable for gene transformation. Still, it has a low tuber root yield, low nutritional quality, high viral and bacterial disease sensitivity, and no sense for farming plants.

Establishing genetic transformation systems for the leading local planting cassava cultivars is of great significance for the genetic improvement of cassava. Genetic transformation protocols for local planting cultivars in Africa and South America have been established, such as TME 204 (the elite East African farmer-preferred cassava cultivar) (Chauhan et al., 2015), T200 (the South African industry-preferred cultivar) (Chetty et al., 2013), TME14 (the landrace commonly grown in West, Central, and East Africa cultivars) (Nyaboga et al., 2015), and Verdinha (the Northeast Brazil cultivar) (Lentz et al., 2018). Cassava is also an important cultivation crop, and the consumption of cassava starch is widespread in China. Therefore, developing effective high-throughput genetic transformation capabilities for popular cassava varieties in China is necessary. South Chinese cassava cultivar 8 (*Manihot esculenta* Crantz cv. SC8) is one of the main cassava local planting cultivars in China. It has excellent characteristics, such as a high yield, high starch content, lodging resistance, and strong adaptability. We established a method for producing somatic embryos (SEs) and friable embryogenic calli (FECs) of cassava SC8 in a previous study (Li et al., 2009; Liu et al., 2019). Based on this, in the present study, the factors that influence transformation efficiency were optimized, such as the density of *Agrobacterium* cells, cocultivation time, FEC humidity, and concentration of acetosyringone (AS). An efficient *Agrobacterium*-mediated gene transformation protocol for cassava SC8 has been developed.

The CRISPR/Cas9-mediated gene editing system is an important tool for crop breeding. However, this technology has rarely been used in cassava breeding and gene function research. First, in 2017, Odipio et al. reported on a case of CRISPR/Cas9-mediated editing of the *MePDS* gene in the genome of TME 204 and cv TM60444 cassava cultivars (Odipio et al., 2017). Five cassava genes (*nCBP-1, nCBP-2, GBSS, PTST1*, and *MePDS*) were knocked out by CRISPR/Cas9 technology; however, the homozygous mutations in these genes in the T_0_ generation were very low (Odipio et al., 2017; Bull et al., 2018; Gomez et al., 2019). Expression of Cas9 under the *A. thaliana YAO* promoter (*pYAO*:hSpCas9 binary vector) has been reported to increase the amount of targeted and homozygous mutations in *A. thaliana* in comparison to Cas9 driven by the Cauliflower mosaic virus (CaMV) 35S promoter (Feng et al., 2018). In this research, we used the *pYAO*:hSpCas9 binary vector to knockout the *MePDS* gene, which improved the efficiency of homozygous mutations in cassava SC8.

## 2. Materials and Methods

### 2.1 Production of Friable Embryogenic Callus (FEC) from Cassava SC8

SE from cassava SC8 was induced according to our previous research (Li et al., 2009). The production of FEC was performed according to the protocol described by Nyaboga et al. (2015) with several modifications (Nyaboga et al., 2015). The mature secondary SEs (coral-shaped) under the microscope were divided into small pieces with a sterile syringe, transferred to Greshoff and Doy (GD) medium (containing 12 mg/L picloram, Table S1), and cultured in the dark at 28 °C. After 14-16 days, the newly formed FECs at the edge of SEs were transferred to fresh GD medium and refreshed every 3 weeks for a maximum of five months.

### 2.2. Vector Constructions

*Agrobacterium* strain LBA4404, harboring the pCAMBIA1304 binary vector, optimized transformation factors. The T-DNA region of plasmid pCAMBIA1304 contains the hygromycin selection marker gene (hpt) and the reporter gene β-glucuronidase (*gusA*) fused with a green fluorescent gene (*GFP*). Both the hpt and gusA-GFP genes are driven by the *CaMV35S* promoter.

The *pYAO*:hSpCas9 binary vector was used to establish the gene editing system for cassava SC8. The Cas9 gene is driven by the *pYAO* promoter, which has been reported to be highly expressed in the embryo sac, embryo, endosperm, and pollen of *Arabidopsis*. The gRNA scaffold is driven by the Arabidopsis *U6-26* promoter. The hygromycin selection marker gene (*hpt*) is driven by the *CaMV35S* promoter.

The phytoene desaturase (*MePDS*, Manes.05G193700) gene was used to quantify gene editing efficiency because its mutation can cause albino seedlings. The target side (GCGTACAAAGCTTCCCAGATAGG) was chosen according to the protocol described by Odipio et al., located in the 13^th^ exon (Odipio et al., 2017). The gRNA sequence was added to the *Bsa* I restriction site linker sequence (Table S2). The synthesized target upstream and downstream primers were diluted with ddH_2_O to a concentration of 10 μM, mixed equally at 98 °C for 3 min, cooled at room temperature, and then placed at 16 °C for 10 min. The *pYAO*:hSpCas9 binary vector was digested with *Bsa* I enzyme at 37 °C for 2 h, and the digested fragments were purified. The annealed primers and the purified vector fragments were ligated by T4 ligase at 16 °C for 6 h, and the ligation product was transformed into *E. coli* DH5α. The positive colonies were identified by sequencing. The correct recombinant plasmid *pYAO*:hSpCas9-MePDS-gRNA was transformed into *Agrobacterium* strain LBA4404 by electroporation.

### 2.3. Preparation of infected Agrobacterium cells and incubation with FEC

*Agrobacterium* LBA4404 harboring pCAMBIA1304 or *pYAO*:hSpCas9-MePDS-gRNA streaked on a YEP plate containing kanamycin 50 mg/L and rifampicin 50 mg/L was cultured overnight. A single colony was inoculated into 1 mL of YEP liquid medium containing the same antibiotics and cultured for 20-24 h at 28 °C at 200 rpm; then, all *Agrobacterium*-cultured solutions were added into 50 mL of fresh YEP liquid medium with the same antibiotics, and culturing continued until an OD_600_ of 0.75 was achieved. The *Agrobacterium*-cultured solutions were centrifuged at 5000 rpm, and the *Agrobacterium* cell precipitates with GD liquid medium were resuspended; this step was repeated once to remove antibiotics completely. Finally, the *Agrobacterium* cells were resuspended in GD liquid medium with a number of AS quantities (50 µM, 100 µM, 150 µM, 200 µM, 250 µM, 300 µM) to a final OD_600_ variety (0.05, 0.25, 0.45, 0.65, 0.85). The *Agrobacterium* cell solution can be used for infection after 60 min at room temperature.

*Agrobacterium* cells were cocultured with FECs according to the following steps. Approximately 1 mL of the three-month-old FECs were transferred to fresh GD liquid medium, fully dispersed with a 5 mL sterile pipette tip, shaken for 30 minutes at 50 rpm and 28 °C, and centrifuged at 1000 rpm. Then the supernatant solution was completely removed. The FEC precipitates were suspended in a 1 mL SCV GD liquid medium. Then 10 mL of the above *Agrobacterium* cells were added and evenly mixed, cocultured for 25 min at 50 rpm 28 °C, and centrifuged for 10 minutes at 1000 rpm at room temperature. The supernatant solution was removed, and the agro-cocultured FEC tissues were transferred onto a nylon filter mesh and then placed on sterilized absorbent filter paper to remove excess bacteria and liquid. The nylon filter mesh with the agroinoculated FECs was transferred onto GD medium with AS to cocultivate for several days (1 d, 3 d, 5 d, 7 d) under dark conditions at 22 °C. The agro-co-cultured FECs, with 1 mL GD liquid medium added to them, were treated as wet FECs and not added as dry FECs. Analyses of AS and *Agrobacterium* cell concentrations were done under dry FECs cocultivate conditions for 3 days at 22 °C

### 2.4. Selection and Regeneration of Transgenic Plants

After coculturing, the FECs infected with *Agrobacterium* were washed 3 times with liquid GD medium containing 500 mg/L carbenicillin, transferred to a new nylon filter with absorbent paper underneath, placed on GD medium containing 250 mg/L carbenicillin, and then incubated for 7 days at 28 °C in the dark. After 7 days, the nylon filter was transferred to fresh GD medium supplemented with 250 mg/L carbenicillin and 8 mg/L hygromycin for 7 days at 28 °C in dark conditions. This step was repeated twice, with the hygromycin content gradually increasing from 15 to 20 mg/L. Afterward, the nylon filter with FEC was transferred onto MSN medium (Table S1) supplemented with 250 mg/L carbenicillin and 20 mg/L hygromycin for 6-8 weeks under a 16/8 h photoperiod at 28 °C, and the medium was refreshed every 2 weeks until the green cotyledons became mature. The mature cotyledons were transferred to shoot-inducting medium (CEM, Table S1) supplemented with 100 mg/L carbenicillin and 10 mg/L hygromycin, and the medium was refreshed every 10-15 days until mature shoots were grown.

The mature shoots were cut and transferred onto MS medium (Table S1) supplemented with 50 mg/L carbenicillin and 10 mg/L hygromycin. Untransformed wild-type cassava seedlings were also cultured on MS medium with 10 mg/L hygromycin as a negative control. Two weeks after culturing, the transgenic shoots developed adventitious roots. In contrast, the nontransgenic shoots did not develop these roots.

### 2.5. Analysis of GUS and GFP Expression

To evaluate the efficiency of cassava transformation using the pCAMBIA1304 binary vector, GUS/GFP coexpressing transgenic tissues (FECs, embryos, cotyledons, and plants) were used for GUS staining or GFP detection. The samples were immersed in GUS reaction buffer (Huayueyang Biotech Co., Ltd., China, GT0391), placed in a vacuum for 4 h, and then incubated overnight at 37 °C. The tissues were washed with 70% ethanol for discoloration, and pictures were taken under an ultradeep field microscope. GFP fluorescence in the transformed tissues (embryos, cotyledons, plants) was visualized using a laser emitter (LUYOR-3415RG).

### 2.6. PCR Analysis

Cassava genomic DNA was extracted from the transformed and wild-type cassava using a plant DNA isolation kit (FOREGENE, CHENGDU). The *gusA* and *GFP* genes inserted in the pCAMBIA1304 vector were used to confirm the transgenic lines by PCR analysis using gene-specific primers (Table S2), which amplified 722 bp fragments for *gusA* and 405 bp fragments for GFP. The Cas9 gene was used for confirmation by PCR analysis using gene-specific primers (Table S2) in the transgenic lines of *pYAO*:hSpCas9-MePDS-gRNA, which amplified 876 bp fragments. Plasmid DNA of *pCAMBIA1304/pYAO*:hSpCas9-MePDS-gRNA and nontransformed plant DNA were used as positive and negative controls, respectively. The standard PCR volume was 50 µL, which consisted of a 200 ng DNA template, 10 mM of each primer, 10 mM dNTP mixture, 10× ExTaq buffer, and 2.5 units of ExTaq DNA polymerase. The reaction conditions for *gusA, GFP*, and *Cas9* were 95 °C for 3 min, 35 cycles of 98 °C for 20 s, 58 °C for 30 s, and 72 °C for 45 s, and a final 10 min extension at 72 °C. PCR products were run on 1% agarose gels with nucleic acid dyes and visualized under a UV transilluminator.

### 2.7. Sanger Sequencing and Hi-TOM Sequencing

A pair of specific primers were designed to amplify a 504 bp fragment, which included the target site of the *MePDS* gene (Table S2). Nontransformed plant DNA was used as a positive control. The PCR system and procedures were the same as those described above. PCR products were Sanger sequenced after detection with a 1.5% agarose gel. Sequences and sequencing peak maps were aligned with the wild-type reference sequence of the *MePDS* gene to characterize CRISPR/Cas9-induced mutations.

Gene editing frequency was analyzed using the Hi-TOM program for high-throughput mutations (Liu et al., 2019). The samples were amplified using target-specific primers (Table S2). According to the Hi-TOM protocol, the first round of PCR products was used as a template for the second round of PCR (barcoding PCR). All products of the second-round PCR were pooled in equimolar amounts in a tube and purified using the OMEGA gel extraction kit. The purified product was sent to Novogene for high-throughput sequencing, and the sequencing data were directly uploaded to the website (http://www.hi-tom.net/hi-tom/).

### 2.8. Statistical Analysis

All the experiments were repeated 3 times, and data for all the parameters were analyzed by variance (ANOVA) using SAS 9.2. Duncan’s new multiple range test was used to detect significant differences between means.

## 3. Results and discussion

### 3.1. Effect of Agrobacterium Cell Concentration on Cassava SC8 Transformation

*Agrobacterium* concentration is an important factor that affects the delivery of T-DNA into plant cells. The infection efficiency for cassava SC8 FEC among six *Agrobacterium* cell densities (OD_600_ values of 0.05, 0.25, 0.45, 0.65, and 0.85) were analyzed. GUS staining of the 30 d-infected FECs and GFP fluorescence in the regenerated cotyledons (under hygromycin selection) were used to evaluate the transformation efficiency. The results showed that the optimal *Agrobacterium* cell concentration for cassava SC8 FEC transformation was an OD_600_ of 0.65 (Fig. 1).

**Fig. 1.**
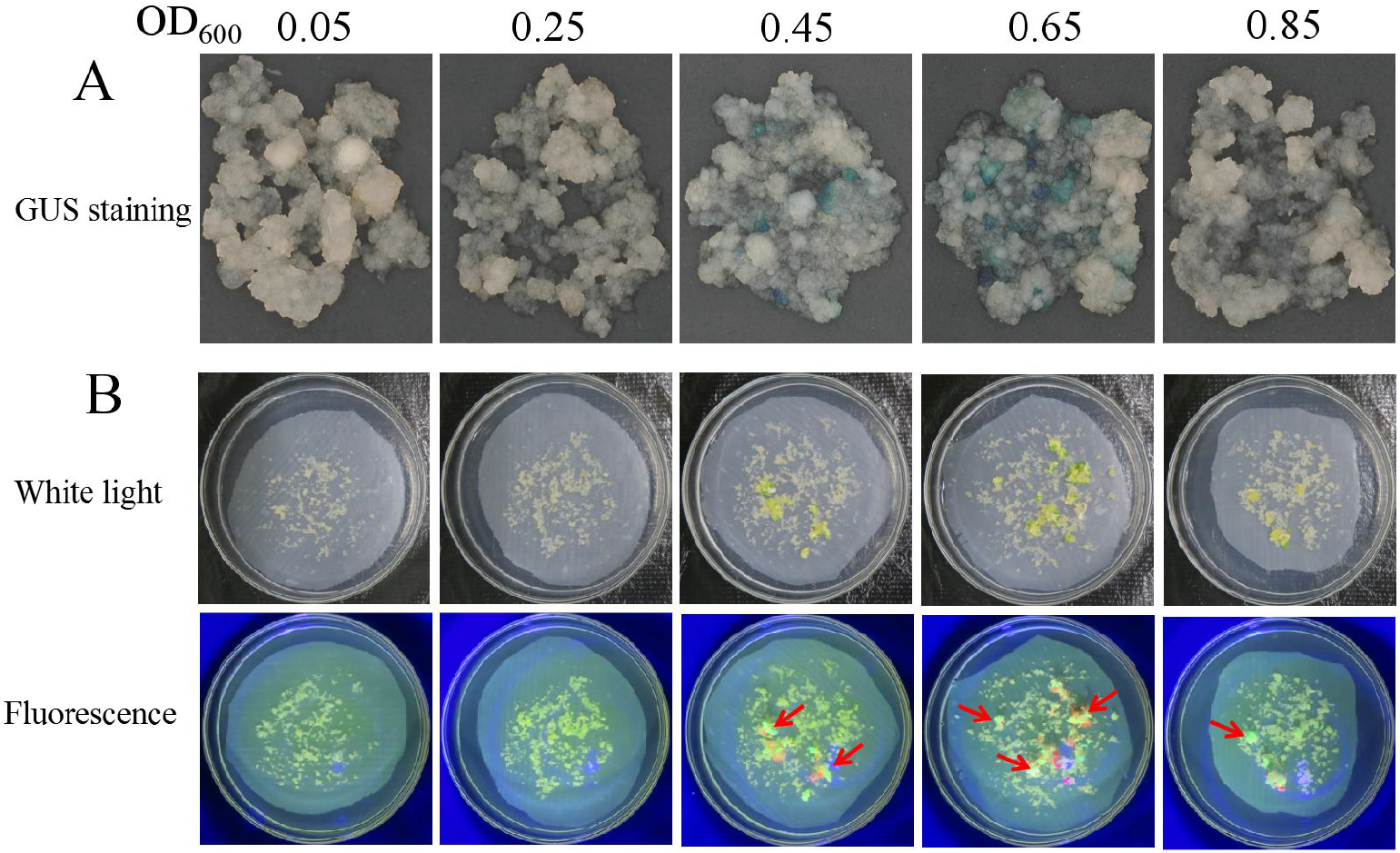
Stable expression of GUS in FECs (A) and GFP in cotyledons (B) under different *Agrobacterium* cell densities. Red arrows point to somatic embryos with green fluorescence.

Furthermore, the effects of the different *Agrobacterium* cell densities on the number of regenerated cotyledons, bud regeneration rate, seedling rate of regenerated plants, rooting rate on the screening medium, PCR positive rate, and several transgenic plants were investigated. The results showed that OD_600_ values of 0.05 and 0.25 did not yield cotyledons. The regeneration rate of buds, seedling rate of regenerated plants, rooting rate on the screening medium, and PCR positive rate were not significantly different under the *Agrobacterium* concentrations of OD_600_ 0.45 and 0.85. In contrast, the *Agrobacterium* concentration of OD_600_ 0.65 produced a significantly (*p*=0.05) higher number of regenerated cotyledons (126.67 ± 41.40) and transgenic plants (88.33 ± 22.55) than other *Agrobacterium* cell densities (Table 1). Based on GUS/GFP expression and the number of regenerated cotyledons and transgenic plants, an *Agrobacterium* concentration OD_600_ of 0.65 was optimal for the genetic transformation of cassava SC8.

**Table 1.**
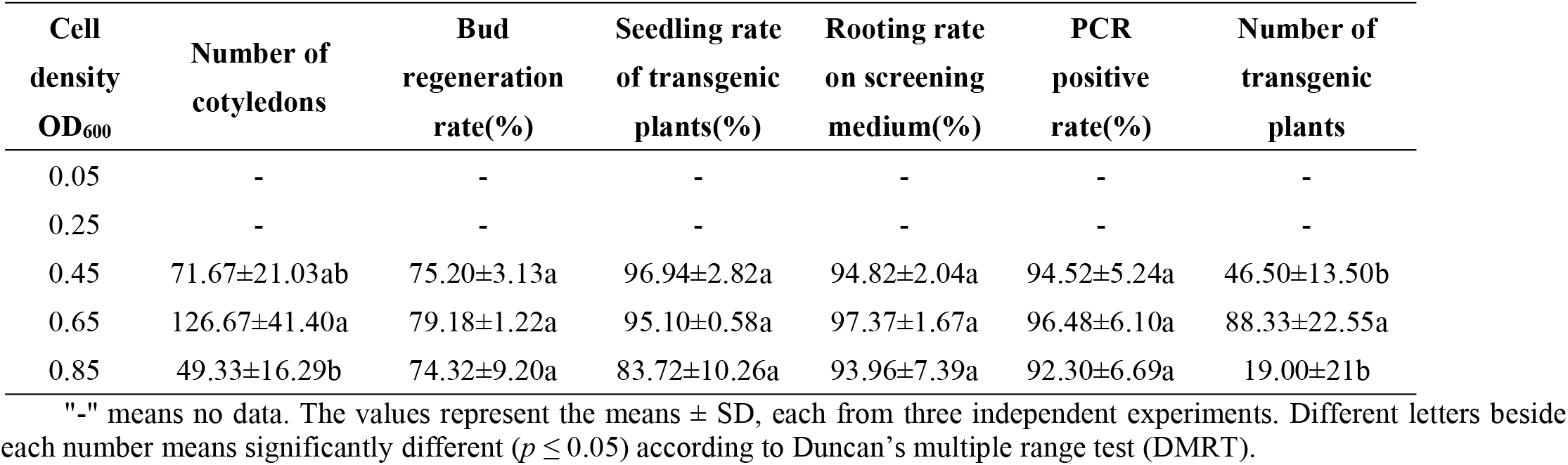
Effect of *Agrobacterium* cell density on cassava SC8 transformation

| Cell density<br>OD<sub>600</sub> | Number of cotyledons | Bud regeneration rate(%) | Seedling rate of transgenic plants(%) | Rooting rate on screening medium(%) | PCR positive rate(%) | Number of transgenic plants |
| --- | --- | --- | --- | --- | --- | --- |
| 0.05 | - | - | - | - | - | - |
| 0.25 | - | - | - | - | - | - |
| 0.45 | 71.67±21.03ab | 75.20±3.13a | 96.94±2.82a | 94.82±2.04a | 94.52±5.24a | 46.50±13.50b |
| 0.65 | 126.67±41.40a | 79.18±1.22a | 95.10±0.58a | 97.37±1.67a | 96.48±6.10a | 88.33±22.55a |
| 0.85 | 49.33±16.29b | 74.32±9.20a | 83.72±10.26a | 93.96±7.39a | 92.30±6.69a | 19.00±21b |

"-" means no data. The values represent the means ± SD, each from three independent experiments. Different letters beside each number means significantly different ( $p \leq 0.05$ ) according to Duncan's multiple range test (DMRT).

### 3.2. Effect of AS Concentration on Cassava SC8 Transformation

Acetosyringone (AS) can induce the efficient expression of the *Vir* gene in the *Agrobacterium* Ti or Ri plasmid and is an important factor for *Agrobacterium*-mediated transformation. Based on the optimal concentration of *Agrobacterium* cells, the effect of AS on the transformation of cassava SC8 was assessed at 50, 100, 150, 200, 250, and 300 µM. GUS staining of the infected FECs after 30 d and GFP fluorescence on the regenerated cotyledons was used to evaluate the transformation efficiency. The results showed that the optimal AS concentration for SC8 FEC transformation was 250 µM, which had stronger GUS staining and GFP fluorescence (Fig. 2) and produced a higher number of regenerated cotyledons (96.33±10.21) and transgenic plants (59.67±4.93) than other AS concentrations (Table 2).

**Fig. 2.**
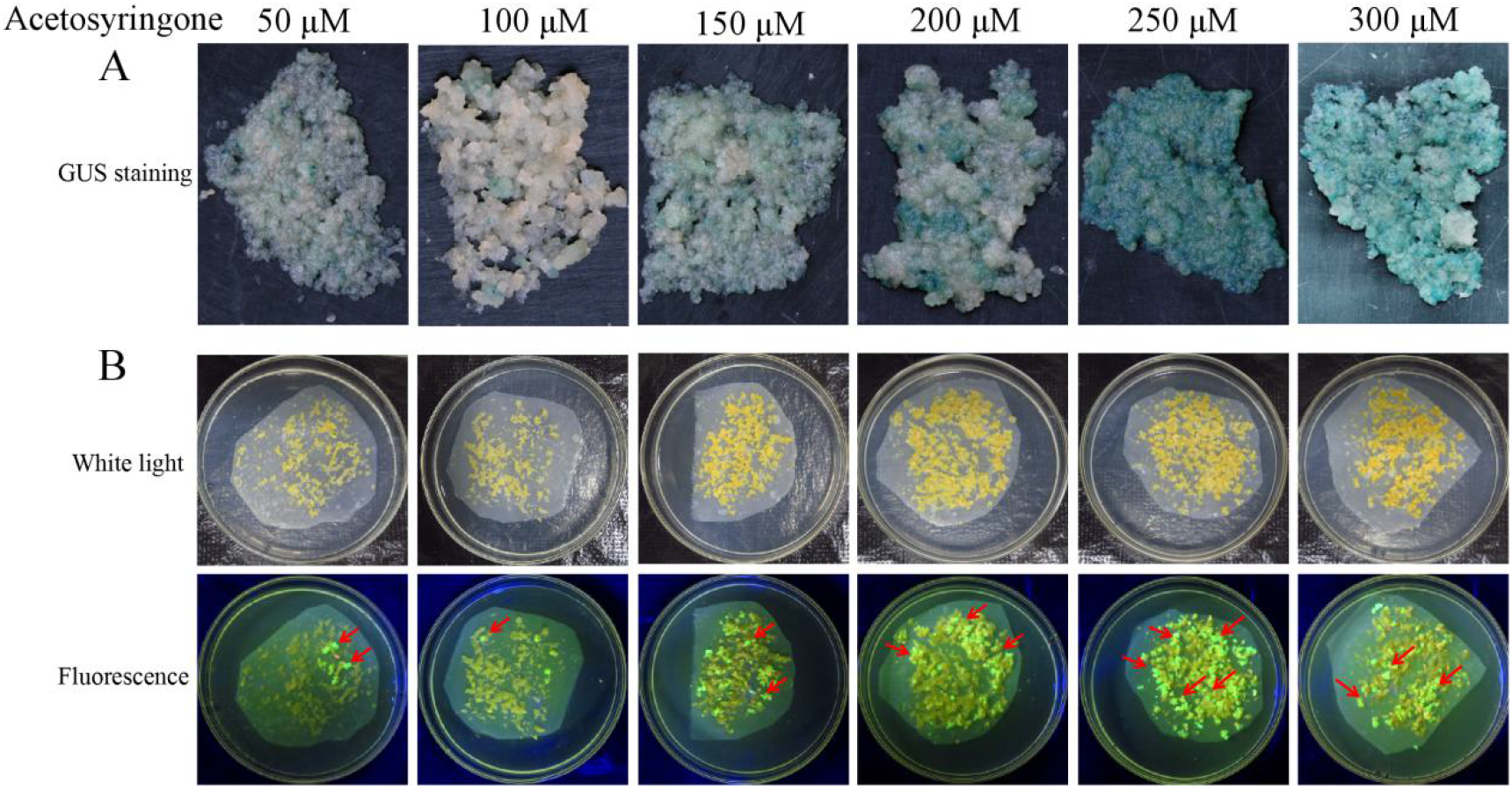
Stable expression of GUS in FECs (A) and GFP in cotyledons (B) under different concentrations of acetosyringone. Red arrows point to somatic embryos with green fluorescence.

**Table 2.** Effect of AS concentration on cassava SC8 transformation

| Treatmen<br>t (μM) | Number of<br>cotyledons | Bud regeneration<br>rate (%) | Seedling rate of<br>transgenic<br>plants (%) | Rooting rate on<br>screening<br>medium (%) | PCR positive<br>rate (%) | Number of<br>transgenic plants |
| --- | --- | --- | --- | --- | --- | --- |
| 50 | 48.00±12.53c | 71.18±10.76ab | 88.42±2.30ab | 88.37±3.63a | 94.20±5.19a | 25.67.±10.02c |
| 100 | 51.67±2.08c | 69.68±4.37ab | 91.50±3.45a | 87.78±1.40a | 93.78±6.01a | 27.00±5.03c |
| 150 | 55.33±5.86bc | 59.96±9.38b | 84.89±1.78b | 89.40±2.84a | 93.76±5.57a | 23.00±1.53c |
| 200 | 70.00±13.23b | 74.04±10.02ab | 86.32±0.81b | 91.03±3.60a | 94.34±4.97a | 38.50±1.53b |
| 250 | 96.33±10.21a | 82.51±3.42a | 89.78±3.94ab | 89.45±3.06a | 93.84±5.39a | 59.67±4.93a |
| 300 | 61.00±6.24bc | 62.58±4.79ab | 86.26±1.61b | 89.06±4.02a | 89.64±1.39a | 28.00±4.16c |

The values represent the means ± SD, each from three independent experiments. Different letters beside each number indicate significant differences ( $p \leq 0.05$ ) according to Duncan's multiple range test (DMRT)

**Fig. 3.**
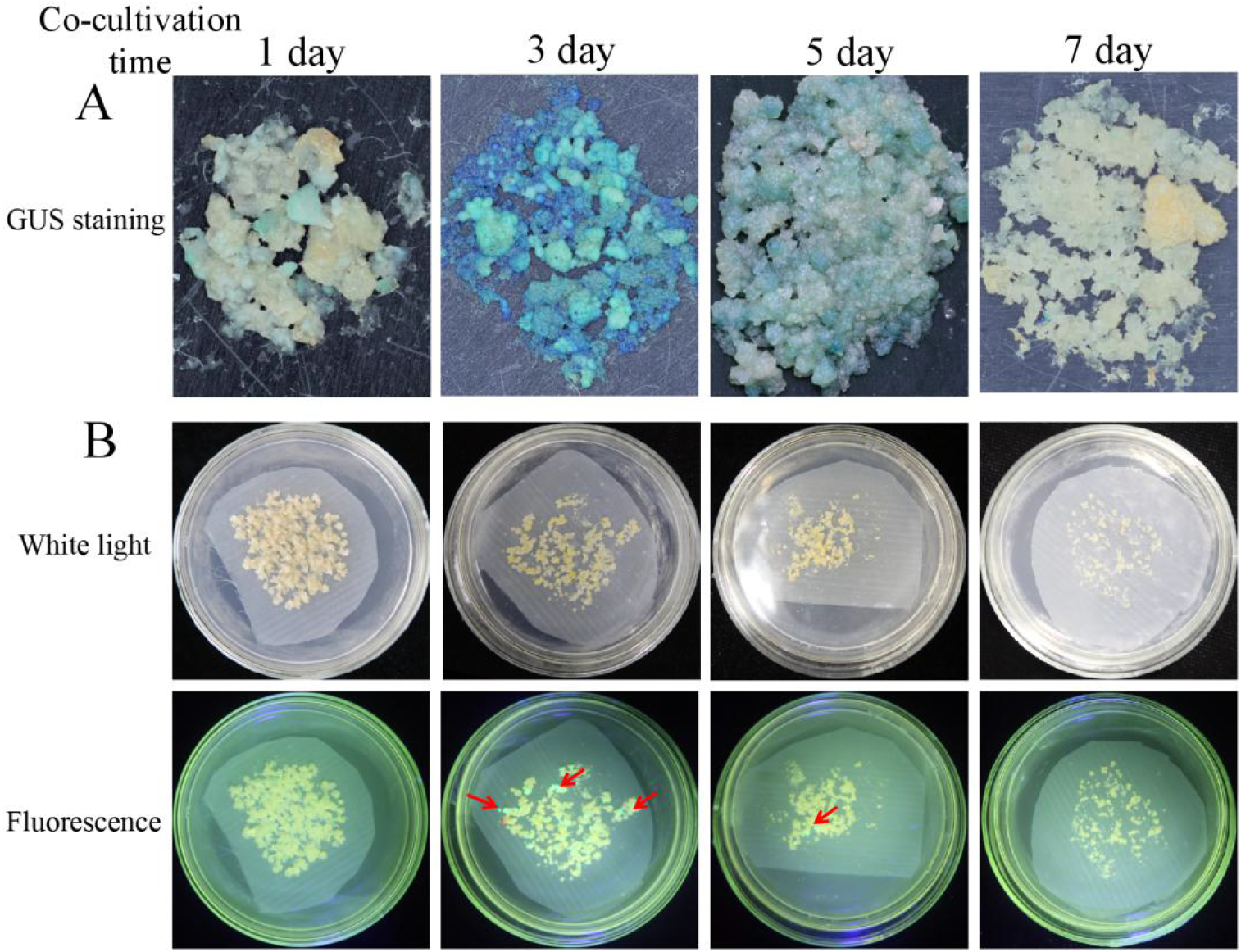
Stable expression of GUS in FECs (A) and GFP in cotyledons (B) under different cocultivation days. Red arrows point to somatic embryos with green fluorescence.

**Fig. 4.**
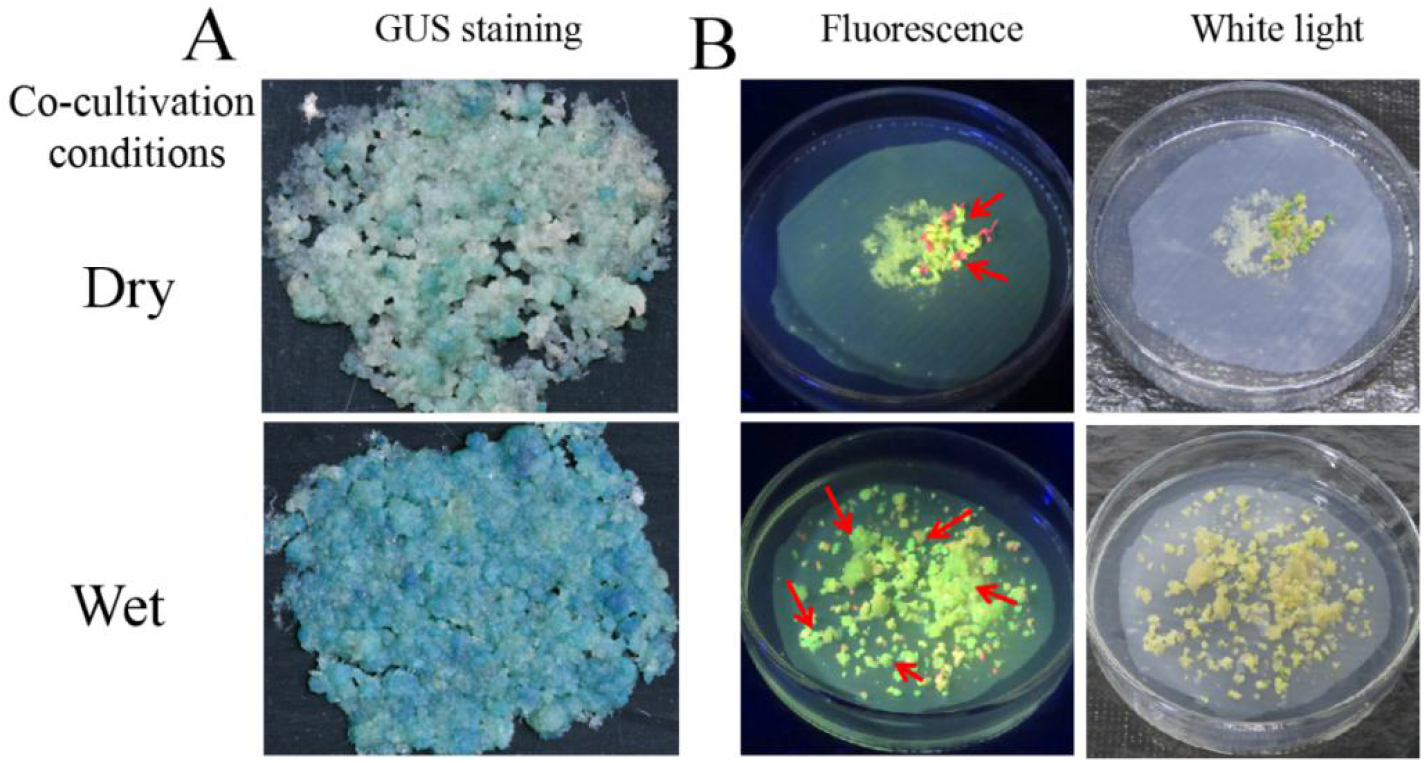
Stable expression of GUS in FECs (A) and GFP in cotyledons (B) under FECs with dry or wet treatment. Red arrows point to somatic embryos with green fluorescence.

### 3.3. Effect of Cocultivation Conditions on Cassava SC8 Transformation

Cocultivation conditions, such as days of cocultivation and the FEC’s humidity, are important factors that need to be optimized in *Agrobacterium*-mediated transformation systems. The results from the GUS staining on the 30 d infected FEC. The GFP fluorescence on the regenerated cotyledons showed that the number of regenerated cotyledons and transgenic plants under the three day cocultivation period were significantly (*p*=0.05) higher than other periods (Table 3); the regenerated cotyledons’ number, bud regeneration rate, seedling rate, rooting rate, and transgenic plant numbers were significantly (*p*=0.05) higher on the wet FEC than on the dry FEC (Table 4). Thus, a three-day cocultivation and wet FEC could increase the transformation efficiency of cassava SC8.

**Table 3.** Effect of cocultivation period on cassava SC8 transformation

| Treatment | Number of cotyledons | Bud regeneration rate(%) | Seedling rate of transgenic plants(%) | Rooting rate on screening medium(%) | PCR positive rate(%) | Number of transgenic plants |
| --- | --- | --- | --- | --- | --- | --- |
| 1 day | - | - | - | - | - | - |
| 3 day | 189.00±33.72<br>a | 75.66±1.62a | 97.00±3.46a | 91.52±7.38a | 94.26±6.63a | 124.00±32.14a |
| 5 day | 83.33±9.07b | 73.46±1.94a | 98.01±3.45a | 94.62±9.10a | 100.00±0a | 55.50±8.14b |
| 7 day | - | - | - | - | - | - |

"-" means no data. The values represent the means ± SD, each from three independent experiments. Different letters beside each number indicate significant differences ( $p \leq 0.05$ ) according to Duncan's multiple range test (DMRT).

**Table 4.** Effect of FEC with dry or wet treatment on cassava SC8 transformation

| Treatment | Number of cotyledons | Bud regeneration rate(%) | Seedling rate of transgenic plants(%) | Rooting rate on screening medium(%) | PCR positive rate(%) | Number of transgenic plants |
| --- | --- | --- | --- | --- | --- | --- |
| dry | 119.67±19.55b | 60.32±3.97b | 85.25±2.17b | 59.50±1.42b | 93.18±6.20a | 34.00±5.57b |
| wet | 201.67±9.50a | 81.11±6.85a | 95.87±1.77a | 84.76±4.89a | 90.81±2.65a | 122.50±17.69a |

The values represent the means ± SD, each from three independent experiments. Different letters beside each number indicate significant differences ( $p \leq 0.05$ ) according to Duncan's multiple range test (DMRT).

### 3.4. Construction of the Optimized Cassava SC8 Transformation Protocol

Axillary bud in vitro cassava SC8 plantlets were cultured on CIM to induce SEs (Fig. 5A-D). The embryos were completely divided and placed on GD to generate sufficient amounts of FEC (Fig. 5E). *Agrobacterium* strain LBA4404 harboring the binary vector pCAMBIA1304 (GUS- and GFP-fused genes driven by the CaMV 35S promoter) was employed for FEC infection, in which *Agrobacterium* strain cell density (OD_600_ of 0.65), 250 µM AS concentration, FEC wet treatment, and a three-day cocultivation period under dark conditions were employed (Fig. 5F). The infected FECs were transformed to MSN medium with hygromycin (8, 15, 20 mg/L) to induce cotyledon initiation (Fig. 5G). Mature cotyledons developed on CEM with 50 mg/L carbenicillin (Fig. 5H). Mature cotyledons were enlarged, and the leaves and shoots were initially developed on COM with 50 mg/L carbenicillin (Fig. 5I). The shoots were placed on MS medium with 50 mg/L carbenicillin to generate transgenic plantlets (Fig. 5J). Roots were induced from the transgenic plantlets on MS with 50 mg/L carbenicillin and 10 mg/L hygromycin (Fig. 5K). After molecular identification, the transgenic plants were transferred to soil (Fig. 5L). Based on the above experimental protocol, three independent gene transformation experiments were tested for transformant efficiency by GUS staining, GFP detection (Fig. 6), and PCR (Fig. S1A). The results showed that approximately 124-143 transgenic lines were generated from 1 mL SCV of agroinfected FEC in approximately five months (Table 5).

**Fig. 5.**
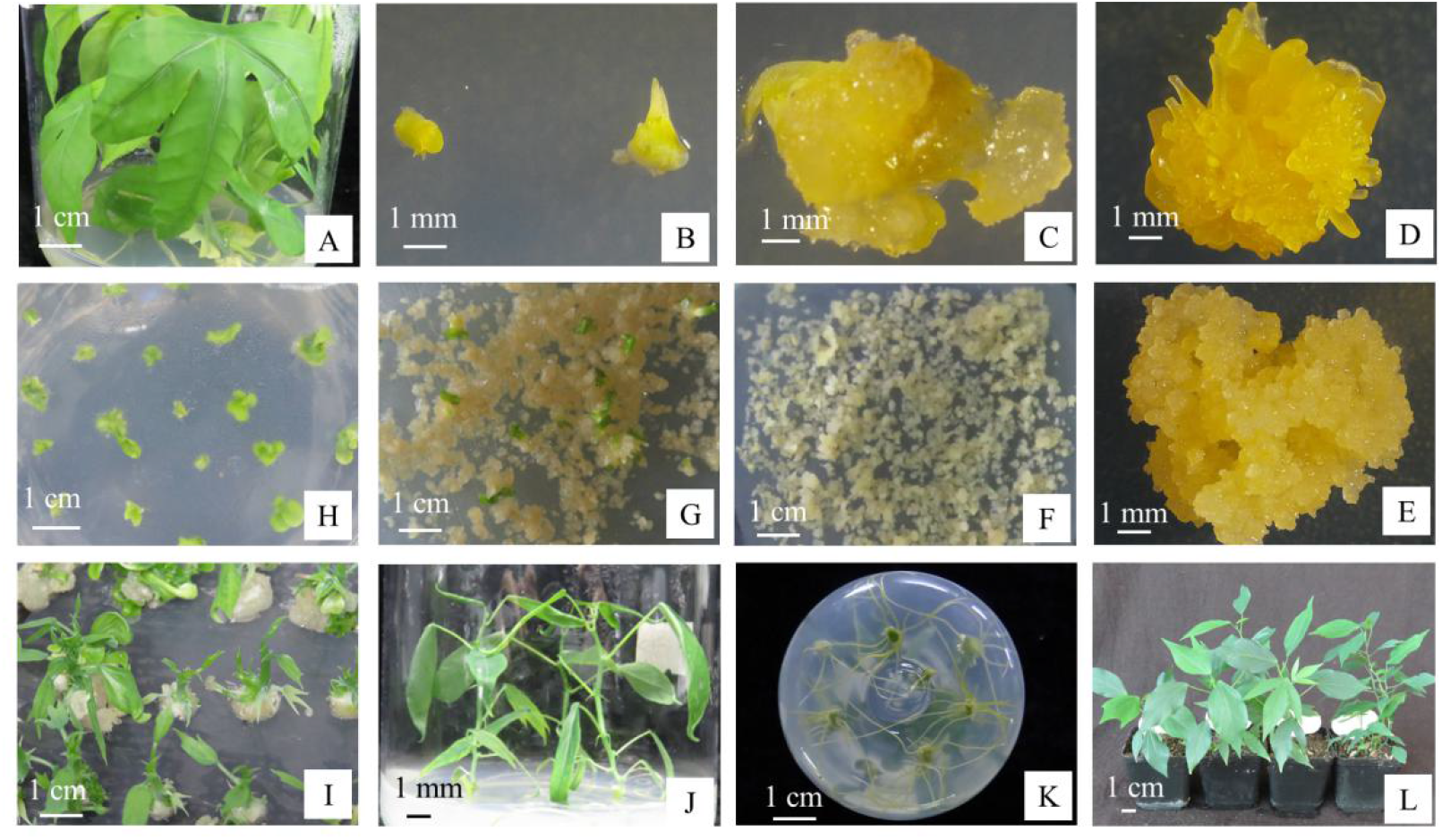
*Agrobacterium*-mediated genetic transformation of cassava SC8 FEC (A). In vitro shoot culture; (B) Axillary bud; (C) Primary SE on CIM medium; (D) SE on CIM medium; (E) Friable embryogenic callus on GD medium; (F) *Agrobacterium*-infected FEC proliferating on GD medium; (G) Developing cotyledons on MSN medium; (H) Cotyledons on CEM medium; (I) Developing shoots on COM medium; (J) Transgenic plantlets on MS medium; (K) Rooting assay of transgenic plants on MS+50 mg/L carbenicillin+10 mg/L hygromycin; (L) Transgenic plants in the soil.

**Table 5.** Validation of the optimized cassava SC8 transformation in three independent experiments by using 1 mL SCV FEC

| Experiments | Number of cotyledons | Number of transgenic plants |
| --- | --- | --- |
| 1 | 187 | 132 |
| 2 | 170 | 124 |
| 3 | 191 | 143 |

**Fig. 6.**
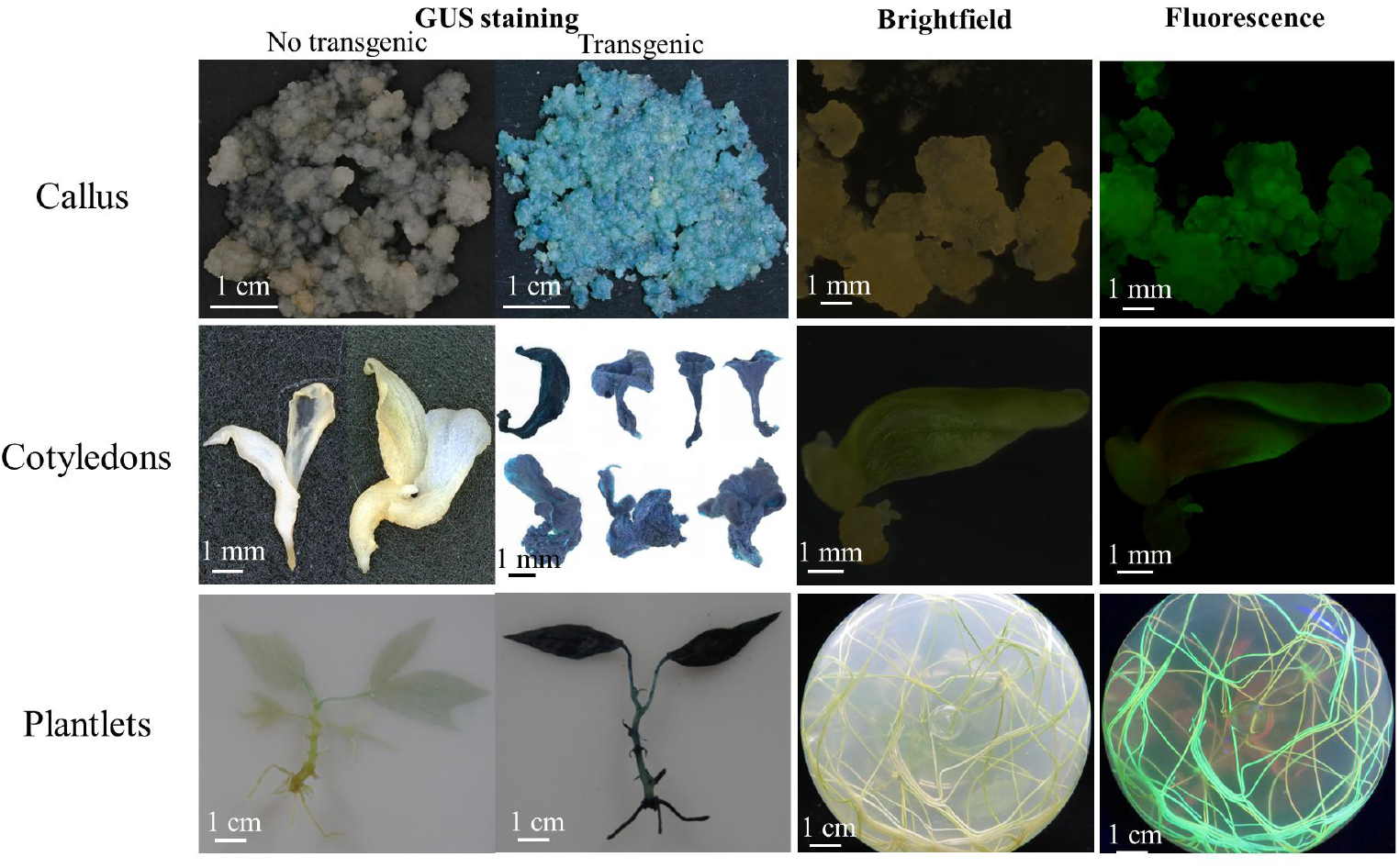
Assessments of the coexpression of GUS and GFP expression in transgenic tissues.

### 3.5. CRISPR/Cas9-Mediated Mutagenesis in Cassava SC8

Using the optimized transformation system, the efficiency of CRISPR/Cas9-mediated gene editing in cassava SC8 was examined. The target side of the *MePDS* gene located in the 13^th^ exon was chosen according to the research of Odipio et al. (Odipio et al., 2017). The *YAO* promoter-driven CRISPR/Cas9 vector *pYAO*:hSpCas9-MePDS-gRNA was constructed to mutate the *MePDS* gene, and the gRNA scaffold was driven by the *Arabidopsis* U6-26 promoter (Fig. 7).

**Fig. 7.**
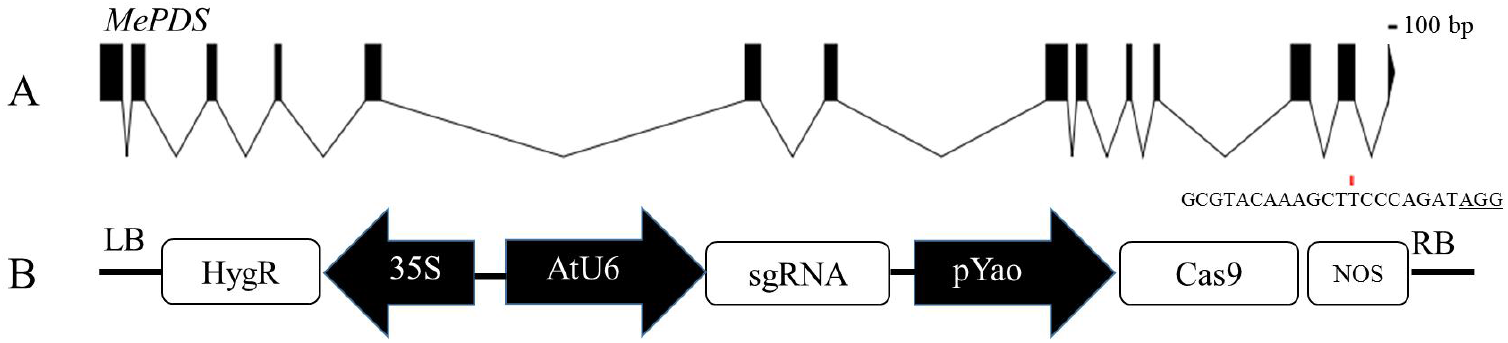
Target site of the *MePDS* gene and the T-DNA of the *pYAO*:hSpCas9-gRNA binary vector. (A) Structural organization of the *MePDS* gene. Exons and introns are shown as boxes and lines, respectively. (B) Schematic of the CRISPR/Cas9 binary vector *pYAO*:hSpCas9-MePDS-gRNA for *MePDS* gene editing through *Agrobacterium*-mediated transformation.

The CRISPR/Cas9 vector with the target sequence was transformed into SC8 FECs using the optimized *Agrobacterium* transformation protocol. A total of 123 independent lines of the regenerated cotyledons were obtained (Fig. 8A). In total, 111 cotyledons were albino (90.24%), 6 cotyledons were yellow or partially albino (4.88%), and 6 cotyledons were green (4.88%) (Fig. 8B).

**Fig. 8.**
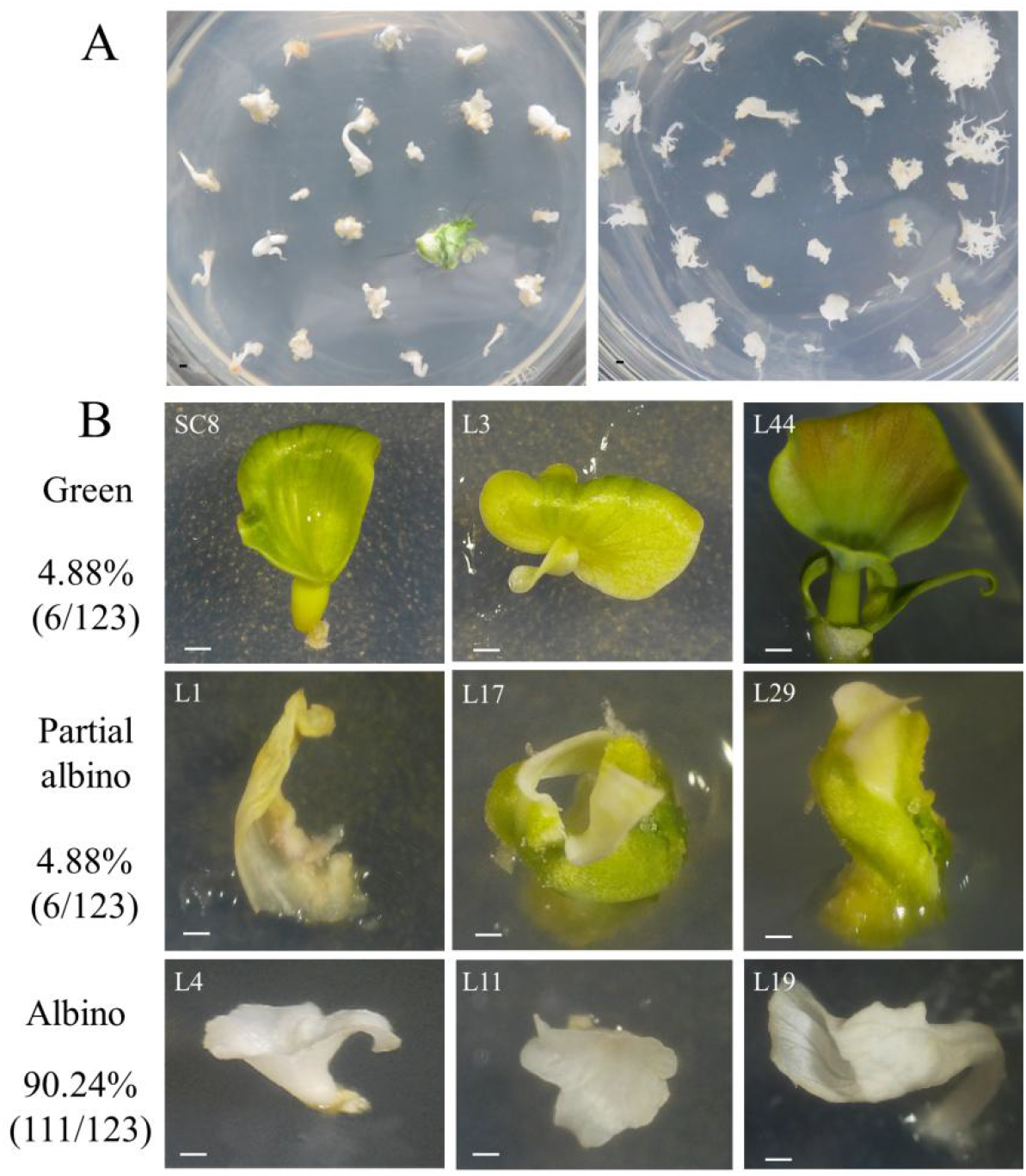
Phenotypes of the regenerated cotyledons after *MePDS* gene editing. (A) Regenerated cotyledons on CEM medium. (B) Phenotypic diversity of CRISPR/Cas9-induced *MePDS* mutations in cassava cotyledons.

A total of 39.02% of the regenerated cotyledon lines successfully germinated plantlets, which generated 48 independent transgenic plant lines (Fig. 9A and Fig. S1B). The target region (250 bp) of the *MePDS* gene from the albino plants was amplified and sequenced by the Sanger method (Fig. 9B and Fig. S1C). The target site of the *MePDS* gene in the L11 and L14 lines showed a double-peak pattern, indicating that the gene editing event occurred in heterozygous types. The peak pattern from Line L4 did not show a double-peak pattern, while a 1 bp insertion at the target site was adjacent to the PAM.

**Fig. 9.**
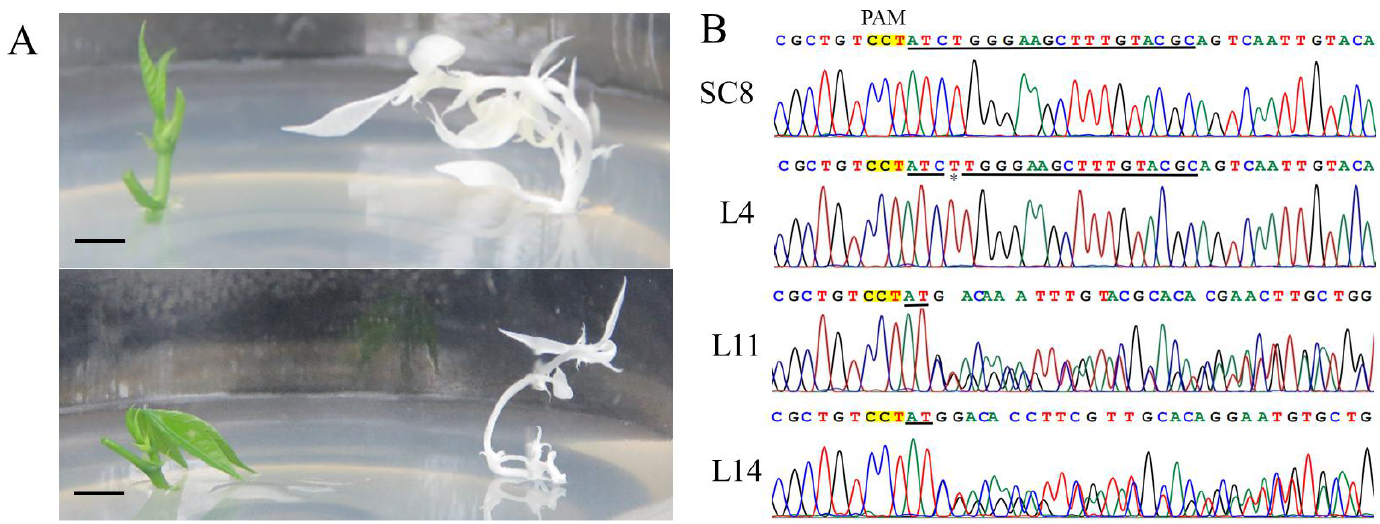
*MePDS* editing of transgenic albino SC8 cassava plants and Sanger sequences. (A) Albino plants and green plants of the *MePDS*-edited transgenic albino SC8 cassava plants. (B) Sanger sequence of the target sites in the *MePDS*-edited transgenic albino SC8 cassava plants.

To further analyze the mutation types of each transgenic line, high-throughput sequencing of the amplified *MePDS* gene target fragments from the 48 transgenic plant lines was performed using Hi-TOM technology (Liu et al., 2019). Most mutations generated by editing were insertions or deletions, which were usually close to the DSB site 3 bp upstream of the PAM (Table S3). In summary, 93.75% of the *MePDS*-edited transgenic cassava SC8 plants had at least one mutation at the target site of the *MePDS* gene, while 6.25% had no mutation (Fig. 8A). Among them, three types of homozygous mono-allelic mutations (45.83%), homozygous bi-allelic mutations (29.16%), and heterozygous mutations (18.75%) were found (Table 6). Notably, using the same target site, the homozygous mutation rate from the *CaMV35S* promoter-driven CRISPR/Cas9 vector in cassava 60444 and TME204 was very low, and the heterozygous mutations were 66.67% -77.78% (Table 6). Thus, these results demonstrated that the *pYAO*:hSpCas9 vector could efficiently create homozygous mutations in cassava.

**Table 6.** Comparison of the mutation types at the same target site by *YAO* or *CaMV35S* promoter-driven CRISPR/Cas9 vector

| Variety | Promoter | Plant lines analyzed | Mutation efficiency | Homozygous mono-allelic | Homozygous bi-allelic | Heterozygous |
| --- | --- | --- | --- | --- | --- | --- |
| SC8 a | *YAO* | 48 | 93.75% (45/48) | 45.83% (22/48) | 29.16% (14/48) | 18.75% (9/14) |
| 60444 b | *CaMV35S* | 9 | 100.00% (9/9) | 11.11% (1/9) | 11.11% (1/9) | 77.78% (7/9) |
| TME204 b | *CaMV35S* | 9 | 100.00% (9/9) | 0.00% (0/9) | 33.33% (3/9) | 66.67% (6/9) |

Note: a, the data from this research; b, the data from Odipio et al. 2017.

### 3.6. Discussion

Since the successful transformation of cassava was first reported in the 1990s, the stress resistance (Xu et al., 2014; Ruan et al., 2017), nutrition (Li et al., 2015; Beyene et al., 2018; Narayanan et al., 2019) and starch properties (Raemakers et al., 2005; Zhao et al., 2011; Ligaba-Osena et al., 2018) of cassava have been improved by transgenic technology. However, these improvements were under the cultivar cv TMS60444 background, which is unsuitable for extensive cassava cultivars. This study established an efficient genetic transformation protocol for cassava SC8, which is the main cultivar in China. An *Agrobacterium* cell density of OD_600_ of 0.65 is best for cassava SC8 genetic transformation. Meanwhile, it has been reported that OD_600_ values of 0.05, 0.25, and 0.50 are suitable for cassava cultivars of TME204 (Chauhan et al., 2015), TME14 (Nyaboga et al., 2015), and TMS60444 (Bull et al., 2009), respectively, which implies that the *Agrobacterium* cell densities for the genetic transformation of different cassava cultivars are different. The addition of AS under cocultivation can significantly improve the transformation efficiency. Six AS concentration gradients (50-300 µM) were set up to optimize the genetic transformation system of cassava SC8.

The results showed that the transgenic efficiency of cassava SC8 was highest when the AS concentration was 250 µM. No research to date has reported on the effects of the concentration of AS on genetic transformation efficiency in different cassava cultivars. In previous studies, 200 µM AS was used for the genetic transformation of farmer-preferred cassava cultivars (such as TME14, TME14, TME204) and the model cultivar cv TMS60444. The FEC of cassava SC8 cannot be fully infected when the cocultivation period is too short (1-2 days); a three-day cocultivation period was found to be best for cassava SC8. The cocultivation periods for other cultivars vary from 2-4 days (Taylor et al., 2012; Chetty et al., 2013). In cassava transformation, the effect of FEC humidity under cocultivation has not been reported in previous studies. Interestingly, FEC wet conditions could increase the transgenic efficiency of cassava SC8. Based on the optimized transformation system, approximately 120-140 transgenic lines per mL SCV were regenerated for cassava SC8 in approximately 5 months. This transformation frequency is significantly higher than previous studies that used model cultivar 60444 and cultivars of TME14 and T200 (Taylor et al., 2012; Nyaboga et al., 2015; Okwuonu et al., 2015).

The CRISPR/Cas9 system has been used for cassava gene editing through a genetic transformation with the *CaMV35S* promoter-driven CRISPR/Cas9 vector (Odipio et al., 2017). The frequency of homozygous mutations is important for cassava mutants. Constitutive promoters usually result in low efficiency of homozygous mutation. It has been reported that cell division promoters (such as the *YAO* and *CDC45* promoters) for CRISPR/Cas9 gene editing could increase the yield of homozygous mutants (Feng et al., 2018). In this research, we used the *pYAO*:hSpCas9 binary vector for the knockout of the *MePDS* gene to assess the homozygous editing rate in cassava SC8. The same target site in the *MePDS* gene has been studied by Odipio et al. using the *CaMV35S* promoter-driven CRISPR/Cas9 vector (Odipio et al., 2017). The expression of Cas9 under the cell division-specific promoter *YAO* produced a mutation rate of 93.75%, which was lower than that of Cas9 driven by the constitutive promoter *CaMV35S* (100.00%). Meanwhile, a high proportion of homozygous mono-allelic mutations (45.83%) were identified from the *YAO* promoter-driven CRISPR/Cas9 vector in cassava SC8, but only 11.11% in TMS60444 and 0.00% in TME204 when Cas9 was driven by the *CaMV35S* promoter (Table 6). It is worth noting that cassava is a highly heterozygous species, and it is difficult to obtain homozygous mutations by hybridization. This study’s high efficiency of homozygous mutations indicates that *YAO* promoter-driven CRISPR/Cas9 could be implemented as a viable approach for cassava genetic improvement.

In conclusion, the efficient genetic transformation and CRISPR/Cas9 gene editing of cassava SC8, one of the main cassava varieties in China, has been reported for the first time in this study. This method was found to efficiently generate 120-140 transgenic lines per mL SCV in approximately 5 months and up to 45.83% homozygous mono-allelic mutations by *YAO* promoter-driven CRISPR/Cas9. These results will be beneficial for the genetic improvement of cassava SC8.

## 4. Conclusions

In this study, an efficient transformation system of cassava SC8 mediated with *Agrobacterium* strain LBA4404 was presented for the first time, in which the factors of *Agrobacterium* strain cell infection (density OD_600_ = 0.65), 250 µM AS induction, and agro-cultivation with wet FEC for three days in dark conditions were found to increase transformation efficiency through the binary vector pCAMBIA1304 harboring GUS- and GFP-fused genes. Based on the optimized transformation protocol, approximately 120-140 independent transgenic lines per mL SCV of FEC by gene transformation in approximately 5 months and 45.83% homozygous mono-allelic mutations of the *MePDS* gene by the *YAO* promoter-driven CRISPR/Cas9 system were generated. This study will open a more functional avenue for the genetic improvement of cassava SC8.

## Credit author statement

**Yajie Wang, Xiaohua Lu**, and **Xinghou Zhen** were responsible for all aspects of the research, including experimental design, data acquisition and analysis, and manuscript preparation. **Hui Yang** and **Yannian Che** worked on the preparation of the studied materials and FEC induction. **Jingyi Hou** worked on PCR analysis. **Ruimei Li** and **Jiao Liu** worked on primer design. **Mengting Geng, Xinwen Hu, Yao Yuan**, and **Jianchun Guo** were responsible for the programs and all experiments, critically revised the manuscript, and provided the final approval of the article.

## Acknowledgments

The authors are grateful to Dr. Qi Xie from the Institute of Genetics and Developmental Biology, Chinese Academy of Sciences, for kindly providing the *pYAO*:hSpCas9 binary vector.

## Appendix A. Supplementary data

Supplementary material related to this article can be found, in the online version, at doi:

