## Supplemental for "Establishment of an efficient transformation and CRISPR/Cas9-mediated gene editing system in Chinese local planting cassava (*Manihot esculenta* Crantz) cultivar SC8"

### Supporting information

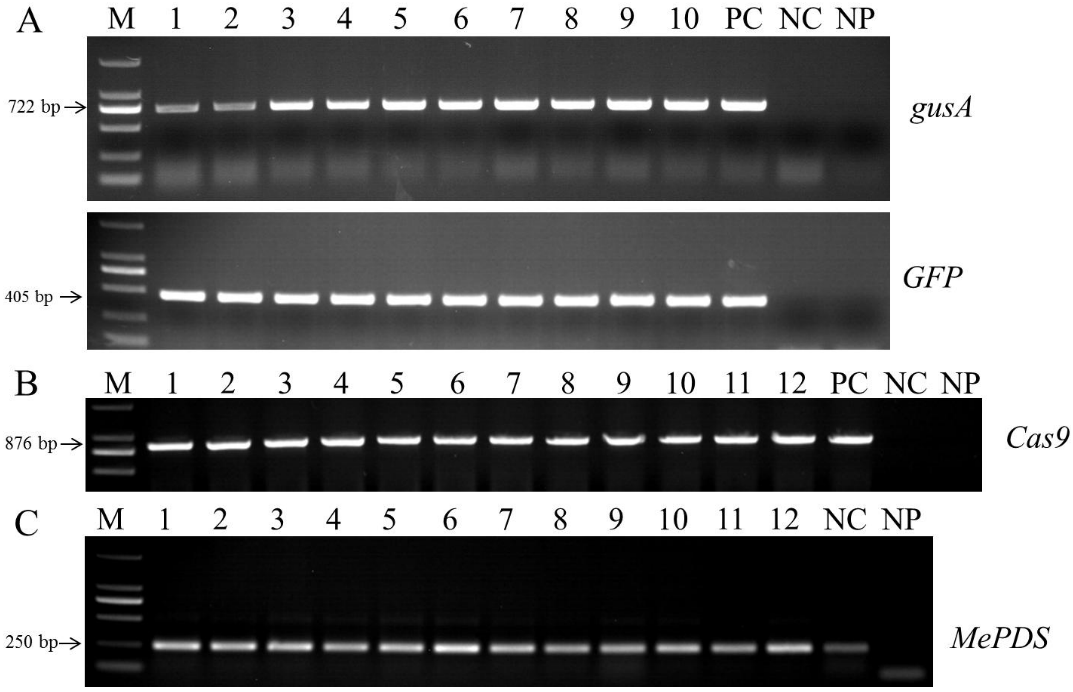


**Supplementary Figure S1. Molecular analysis of transgenic plants.** (A) PCR analysis of transgenic lines of cassava using *gusA* and *GFP* gene specific primers. Lanes M: 2000 bp marker (TAKARA), Lanes 1-10: different transgenic lines, PC: Positive control, NC: Negative control, NP: No template PCR. (B) PCR analysis of CRISPR/Cas9 T-DNA integration. Lanes M: 2000 bp marker (TAKARA), Lanes 1-12: different transgenic lines, PC: Positive control, NC: Negative control, NP: No template PCR. (C) PCR amplification of the *MePDS* target sequence. Lanes M: 2000 bp marker(TAKARA), Lanes 1-12: different transgenic lines, NC: Negative control, NP: No template PCR.

**Supplementary Table S1. The media and their compositions**

| Medium | Compositions (1 L) |
|---|---|
| MS | 4.4 g MS，30 g sucrose，0.32 mg $CuSO_4$，pH5.8，8 g Agar |
| GD | 2.75 g GD，20 g sucrose，12 mg picloram，pH5.8，8 g Agar |
| CIM | 4.4 g MS，20 g sucrose，0.32 mg $CuSO_4$，12 mg picloram，pH5.8，8 g Agar |
| CEM | 4.4 g MS，30 g sucrose，0.32 mg $CuSO_4$，0.4 mg 6-BA，pH5.8，8 g Agar |
| COM | 4.4 g MS，30 g sucrose，0.32 mg $CuSO_4$，1 mg 6-BA，pH5.8，8 g Agar |
| MSN | 4.4 g MS，20 g sucrose，0.32 mg $CuSO_4$，1 mg NAA，pH5.8，8 g Agar |

All the above culture media need to be autoclaved at 121°C for 20 min.

**Supplementary Table S2. Primers used in this study.**

| Primer name | Primer sequence | length | Aims |
|---|---|---|---|
| MePDS-F:<br>MePDS-R: | 5´-AGCTGGGGACTACACAAAGC-3´<br>5´-CCACACCCATTAGGCCTTGTAA-3´ | 210 bp | Amplification of the *MePDS* part |
| GFP-F:<br>GFP-R: | 5´-CAAGGACGACGGCAACTACA-3´<br>5´-TCGTCCATGCCGAGAGTGAT-3´ | 405 bp | Amplification of the *GFP* part |
| GUS-F:<br>GUS-R: | 5´-CCTCGCATTACCCTTACGCT-3´<br>5´-TTTCTTGTTACCGCCAACGC-3´ | 722 bp | Amplification of the *GUSA* part |
| CAS9-F:<br>CAS9-R: | 5´-GCAAGCTGCTCTAGCCAATACGC-3´<br>5´-CGGGAAACGACAATCTGATCCAAG-3´ | 876 bp | Amplification of the Cas9 part |
| MePDS-gRNA-F<br>MePDS-gRNA-R | 5´-gattGCGTACAAAGCTTCCCAGAT-3´<br>5´-aaacATCTGGGAAGCTTTGTACGC-3´ | | *MePDS* gene target |
| MePDS-HT-F:<br>MePDS-HT-R: | 5´-ggagtgagtacggtgtgcAGCTGGGGACTACACAAAGC-3´<br>5´-gagttggatgctggatggCCACACCCATTAGGCCTTGTAA-3´ | 250 bp | *MePDS* gene editing test |

**Supplementary Table S3. Results of *MePDS* gene by Hi-TOM analysis**

| Lines | Sequence near the target | Reads | Ratio | Variation type | Variation sequence |
|---|---|---|---|---|---|
| **WT** | GTCCTATCTGGGAAGCTTTGTACGCAG | 10974 | 96.26% | | |
| **L1** | GTCCTATCTTGGGAAGCTTTGTACGCAG | 766 | 90.22% | 1I | T |
| | GTCCTATCTGGGAAGCTTTGTACGCAG | 83 | 9.78% | WT | |
| **L2** | GTCCTATCTTGGGAAGCTTTGTACGCAG | 6636 | 93.62% | 1I | T |
| **L3** | GTCCTATCTGGGAAGCTTTGTACGCAG | 6588 | 55.67% | WT | |
| | GTCCTATCTTGGGAAGCTTTGTACGCAG | 5100 | 43.09% | 1I | T |
| **L4** | GTCCTATCTTGGGAAGCTTTGTACGCAG | 14197 | 96.76% | 1I | T |
| **L5** | GTCCTATCTTGGGAAGCTTTGTACGCAG | 13236 | 97.92% | 1I | T |
| **L6** | GTCCTATCTTGGGAAGCTTTGTACGCAG | 24680 | 84.58% | 1I | T |
| | | 1489 | 5.10% | 1I,SNP | T,A->G |
| **L7** | GTCCTAT------------------------GTACGCAG | 11295 | 53.29% | 12D | TCTGGGAAGCTT |
|  | GTCCTATC----GGAAGCTTTGTACGCAG | 8993 | 42.43% | 2D | TG |
| **L8** | GTCCTATCTTGGGAAGCTTTGTACGCAG | 14194 | 96.76% | 1I | T |
| **L9** | GTCCTATCTTGGGAAGCTTTGTACGCAG | 12625 | 98.36% | 1I | T |
| **L10** | GTCCTATCTTGGGAAGCTTTGTACGCAG | 19092 | 94.53% | 1I | T |
| **L11** | GTCCTAT------------------------GTACGCAG | 13658 | 52.02% | 12D | TCTGGGAAGCTT |
| | GTCCTATC---GGAAGCTTTGTACGCAG | 11569 | 44.07% | 2D | TG |
| **L12** | GTCCTATCTTGGGAAGCTTTGTACGCAG | 11196 | 50.68% | 1I | T |
|  | GTCCTATC----------------TTTGTACGCAG | 5433 | 24.59% | 8D | CTGGGAAG |
|  | GTCCTATC------------GCTTTGTACGCAG | 5138 | 23.26% | 6D | TGGGAA |
| **L13** | GTCCTATCTTGGGAAGCTTTGTACGCAG | 31334 | 77.67% | 1I | T |
| | | 2770 | 6.87% | 1I,SNP | T,A->G |
| **L14** | GTCCTAT------------------------GTACGCAG | 185 | 29.60% | 12D | TCTGGGAAGCTT |
| | GTCCTATCTTGGGAAGCTTTGTACGCAG | 183 | 29.28% | 1I | T |
|  | GTCCTATC----GGAAGCTTTGTACGCAG | 174 | 27.84% | 2D | TG |
| | GTCCTATCTGGGAAGCTTTGTACGCAG | 83 | 13.28% | WT | |
| **L15** | GTCCTATCTTGGGAAGCTTTGTACGCAG | 6786 | 96.57% | 1I | T |
| **L16** | GTCCTATCTTGGGAAGCTTTGTACGCAG | 495 | 82.91% | 1I | T |
| | GTCCTATCTGGGAAGCTTTGTACGCAG | 102 | 17.09% | WT | |
| **L17** | GTCCTATCTTGGGAAGCTTTGTACGCAG | 142 | 62.01% | 1I | T |
| | GTCCTATCTGGGAAGCTTTGTACGCAG | 87 | 37.99 | WT | |
| **L18** | GTCCTAT------------------------GTACGCAG | 14009 | 53.47% | 12D | TCTGGGAAGCTT |
| | GTCCTATC---GGAAGCTTTGTACGCAG | 11293 | 43.10% | 2D | TG |
| **L19** | GTCCTATCTTGGGAAGCTTTGTACGCAG | 12248 | 75.17% | 1I | T |
|  | GTCCTAT------------------------GTACGCAG | 2099 | 12.88% | 12D | TCTGGGAAGCTT |
|  | GTCCTATC----GGAAGCTTTGTACGCAG | 1665 | 10.22% | 2D | TG |
| **L20** | GTCCTATCTTGGGAAGCTTTGTACGCAG | 13550 | 96.58% | 1I | T |
| **L21** | GTCCTATCTTGGGAAGCTTTGTACGCAG | 10071 | 97.79% | 1I | T |

| | | | | | |
|---|---|---|---|---|---|
| **L22** | GTCCTATCTTGGGAAGCTTTGTACGCAG | 8084 | 85.56% | 1I | T |
| **L23** | GTCCTATCTTGGGAAGCTTTGTACGCAG | 20220 | 87.32% | 1I | T |
| **L24** | GTCCTATC----------------TTTGTACGCAG | 14337 | 47.83% | 8D | CTGGGAAG |
|  | GTCCTATC------------GCTTTGTACGCAG | 13993 | 46.68% | 6D | TGGGAA |
| **L25** | GTCCTATCTTGGGAAGCTTTGTACGCAG | 26563 | 85.38% | 1I | T |
| **L26** | GTCCTAT------------------------GTACGCAG | 14442 | 51.77% | 12D | TCTGGGAAGCTT |
|  | GTCCTATC----GGAAGCTTTGTACGCAG | 12552 | 45.00% | 2D | TG |
| **L27** | GTCCTATCTTGGGAAGCTTTGTACGCAG | 25825 | 92.07% | 1I | T |
| **L28** | GTCCTATCTTGGGAAGCTTTGTACGCAG | 23355 | 93.31% | 1I | T |
| **L29** | GTCCTATCTTGGGAAGCTTTGTACGCAG | 221 | 43.16% | 1I | T |
| | GTCCTATCTGGGAAGCTTTGTACGCAG | 140 | 27.34% | WT | |
|  | GTCCTATC------------GCTTTGTACGCAG | 77 | 15.04% | 6D | TGGGAA |
|  | GTCCTATC----------------TTTGTACGCAG | 74 | 14.45% | 8D | CTGGGAAG |
| **L30** | GTCCTATCTTGGGAAGCTTTGTACGCAG | 27513 | 85.43% | 1I | T |
| **L31** | GTCCTATCTTGGGAAGCTTTGTACGCAG | 20529 | 90.14% | 1I | T |
| **L32** | GTCCTATCTTGGGAAGCTTTGTACGCAG | 19672 | 93.69% | 1I | T |
| **L33** | GTCCTAT------------------------GTACGCAG | 6274 | 51.74% | 12D | TCTGGGAAGCTT |
|  | GTCCTATC----GGAAGCTTTGTACGCAG | 5246 | 43.27% | 2D | TG |
| **L34** | GTCCTATC----------------TTTGTACGCAG | 9147 | 50.13% | 8D | CTGGGAAG |
|  | GTCCTATC------------GCTTTGTACGCAG | 8709 | 47.73% | 6D | TGGGAA |
| **L35** | GTCCTAT------------------------GTACGCAG | 12936 | 52.24% | 12D | TCTGGGAAGCTT |
| | GTCCTATC---GGAAGCTTTGTACGCAG | 11159 | 45.06% | 2D | TG |
| **L36** | GTCCTAT------------------------GTACGCAG | 8962 | 52.69% | 12D | TCTGGGAAGCTT |
|  | GTCCTATC----GGAAGCTTTGTACGCAG | 7243 | 42.59% | 2D | TG |
| **L37** | GTCCTATCTTGGGAAGCTTTGTACGCAG | 28092 | 80.70% | 1I | T |
| | | 2154 | 6.19% | 1I,SNP | T,A->G |
| **L38** | GTCCTAT------------------------GTACGCAG | 11409 | 46.01% | 12D | TCTGGGAAGCTT |
| | GTCCTATC---GGAAGCTTTGTACGCAG | 8189 | 33.03% | 2D | TG |
| | GTCCTATCTTGGGAAGCTTTGTACGCAG | 5029 | 20.28% | 1I | T |
| **L39** | GTCCTATCTGGGAAGCTTTGTACGCAG | 23432 | 90.57% | WT | |
| **L40** | GTCCTATCTTGGGAAGCTTTGTACGCAG | 21323 | 89.51% | 1I | T |
| **L41** | GTCCTATC------------GCTTTGTACGCAG | 9595 | 50.00% | 6D | TGGGAA |
|  | GTCCTATC----------------TTTGTACGCAG | 9308 | 48.50% | 8D | CTGGGAAG |
| **L42** | GTCCTATCTTGGGAAGCTTTGTACGCAG | 24741 | 85.13% | 1I | T |
| **L43** | GTCCTATC--GGGAAGCTTTGTACGCAG | 10392 | 49.40% | 1D | T |
| | GTCCTATCTTGGGAAGCTTTGTACGCAG | 10150 | 48.25% | 1I | T |
| **L44** | GTCCTATCTGGGAAGCTTTGTACGCAG | 18162 | 91.12% | SNP | T->A |
| **L45** | GTCCTATCTTGGGAAGCTTTGTACGCAG | 281 | 49.13% | 1I | T |
| | GTCCTATCTGGGAAGCTTTGTACGCAG | 135 | 23.60% | WT | |
|  | GTCCTATC----GGAAGCTTTGTACGCAG | 78 | 13.64% | 2D | TG |
|  | GTCCTAT------------------------GTACGCAG | 78 | 13.64% | 12D | TCTGGGAAGCTT |
| **L46** | GTCCTATCTTGGGAAGCTTTGTACGCAG | 144 | 53.33% | 1I | T |

| | | | | | |
|---|---|---|---|---|---|
| | GTCCTATCTGGGAAGCTTTGTACGCAG | 126 | 46.67% | WT | |
| **L47** | GTCCTATCTTGGGAAGCTTTGTACGCAG | 373 | 53.52% | 1I | T |
| | GTCCTATCTGGGAAGCTTTGTACGCAG | 324 | 46.48% | WT | |
| **L48** | GTCCTATCTGGGAAGCTTTGTACGCAG | 11013 | 98.55% | WT | |

The yellow areas are the PAM areas, the underlines represent the target sequences; The red bases represent the inserted bases, -- represent the deleted bases, Lowercase letters stand for substitution bases; I: insertion, D: deletion, SNP:substitution, WT: wild type.
